# ProFuMCell and ProModb: web services for analyzing interaction-based functionally localized protein modules in a cell

**DOI:** 10.1101/2021.06.03.446966

**Authors:** Barnali Das, Pralay Mitra

## Abstract

The modular organization of a cell which can be determined by its interaction network allows us to understand a mesh of cooperation among the functional modules. Therefore, cellular level identification of functional modules aids in understanding the functional and structural characteristics of the biological network of a cell and also assists in determining or comprehending the evolutionary signal. We develop ProFuMCell that performs real-time web scraping for generating clusters of the cellular level functional units of an organism. ProFuMCell constructs the Protein Locality Graphs and clusters the cellular level functional units of an organism by utilizing experimentally verified data from various online sources. Also, we develop ProModb, a database service that houses precomputed whole-cell protein-protein interaction network-based functional modules of an organism using ProFuMCell. Our web service is entirely synchronized with the KEGG pathway database and allows users to generate spatially localized protein modules for any organism belonging to the KEGG genome using its real-time web scraping characteristics. Hence, the server will host as many organisms as is maintained by the KEGG database. Our web services provide the users a comprehensive and integrated tool for an efficient browsing and extraction of the spatial locality-based protein locality graph and the functional modules constructed by gathering experimental data from several interaction databases and pathway maps. We believe that our web services will be beneficial in pharmacological research, where a novel research domain called modular pharmacology has initiated the study on the diagnosis, prevention, and treatment of deadly diseases using functional modules.

**Graphical TOC Entry:** 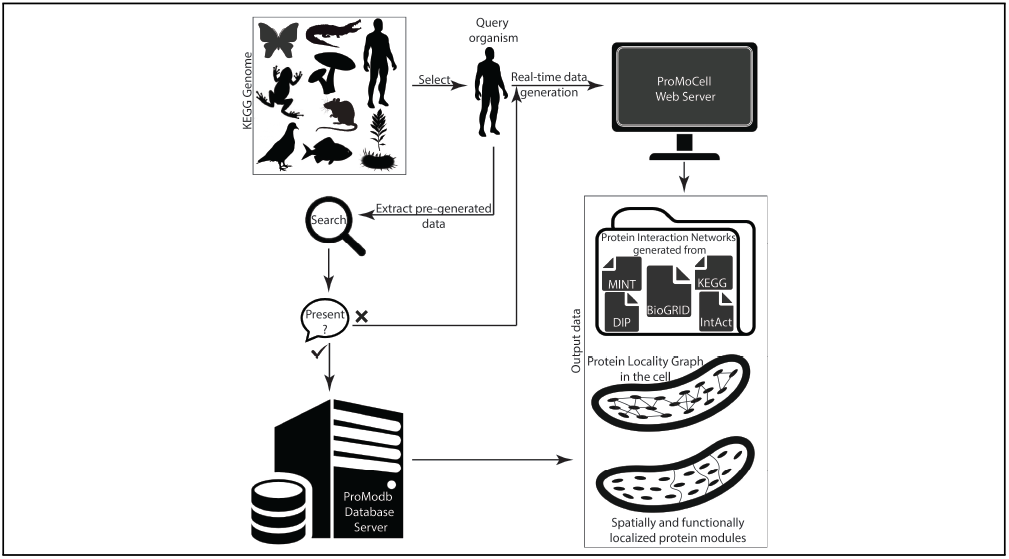

## Introduction

The biological network of a cell exhibits a modular organization where the cellular functions are accomplished by a mesh of cooperating functional modules that comprises of different groups of interacting molecules. Identifying functional modules of a cell is highly crucial as it helps in understanding the functional and structural characteristics of a biological network^1^ and also assists in determining or comprehending the evolutionary signal.^2^ Functional modules play a vital role in the functional organization of a cell, and in the underlying cellular mechanisms.^3^ Barabasi et al.^4^ noted that interruption of the functional modules gives rise to diseases. Whereas, Yu et al. studied the consequences of reorganization of the intra-module or the inter-module connections or interactions on the cellular properties or on the cellular functions. ^2^ Literature evidence informs that proper knowledge of the functional modules can assist in the diagnosis, prevention, and treatment of deadly diseases like cancer.^5,6^ To address the issue, a unique area called modular pharmacology (MP)^7^ has been instigated as a part of the pharmacological research. Therefore, identification of the functional modules is highly crucial and can be beneficial for future research in various domains.

Here, we introduce a web service viz., ProFuMCell (*Pro*tein interaction-based *Fu*nctional *M*odules of the *Cell* of an organism) for constructing the Protein Locality Graphs and clustering the cellular level functional units of an organism by utilizing experimentally verified data from various online sources. ProFuMCell, established on network-based zoning method,^8^ is entirely synchronized with the Kyoto Encyclopedia of Genes and Genomes (KEGG) pathway database^10^ and allows users to generate spatially localized protein modules for any organism belonging to the KEGG genome using its real-time web scraping characteristics. Hence, the server will host as many organisms as is maintained by KEGG database. Currently, ProFuMCell supports 6770 organisms, including 536 eukaryotes, 5577 bacteria, 316 archaea, and 341 viruses. Usually, ProFuMCell takes a few hours of execution time for an organism. Therefore, to facilitate its utility, we develop ProModb (Protein interaction-based functional Module database) that stores precomputed ProFuMCell outputs of different organisms. Presumably, no other services like ProFuMCell or ProModb exist till date.

## Materials and Methods

### The procedure

ProFuMCell implements a recently published network-based zoning approach where spatial localization of the protein molecules in a cell leads towards their functional localization.^8^ It initially constructs the PPI network for the whole-cell utilizing protein-protein interaction (PPI) data from KEGG pathway maps^10^ and PPI databases like Database of Interacting Proteins (DIP),^11^ IntAct,^12^ Molecular Interaction database (MINT),^13^ and BioGRID,^14^ which are termed as GPG, PPID, PPII, PPIM, and PPIB respectively. For each interaction network, ProFuMCell computes protein spatial locality based on the fact that two interacting proteins must be in proximity to each other. Hence, edge weights representing confidence scores of closeness are assigned to the interactions of the PPI network accordingly. Then the server combines all the interaction networks into a single graph viz., the Protein Locality Graph (PLG). Finally, ProFuMCell identifies the functional modules of spatially localized interacting proteins by clustering the PLG (further details are provided at the Supplementary Section S3). For the unicellular bacterium *Escherichia coli K12* (*E. coli*), we demonstrated^8^ that the clustered modules are well within the functional proximity as per the Gene Ontology database.^15^ Figure 1 demonstrates the entire methodology of the ProFuMCell server when input is organism *E. coli*. Further details of the ProFuMCell server methodology are provided in SI Sections S1-S5.

**Figure 1:**
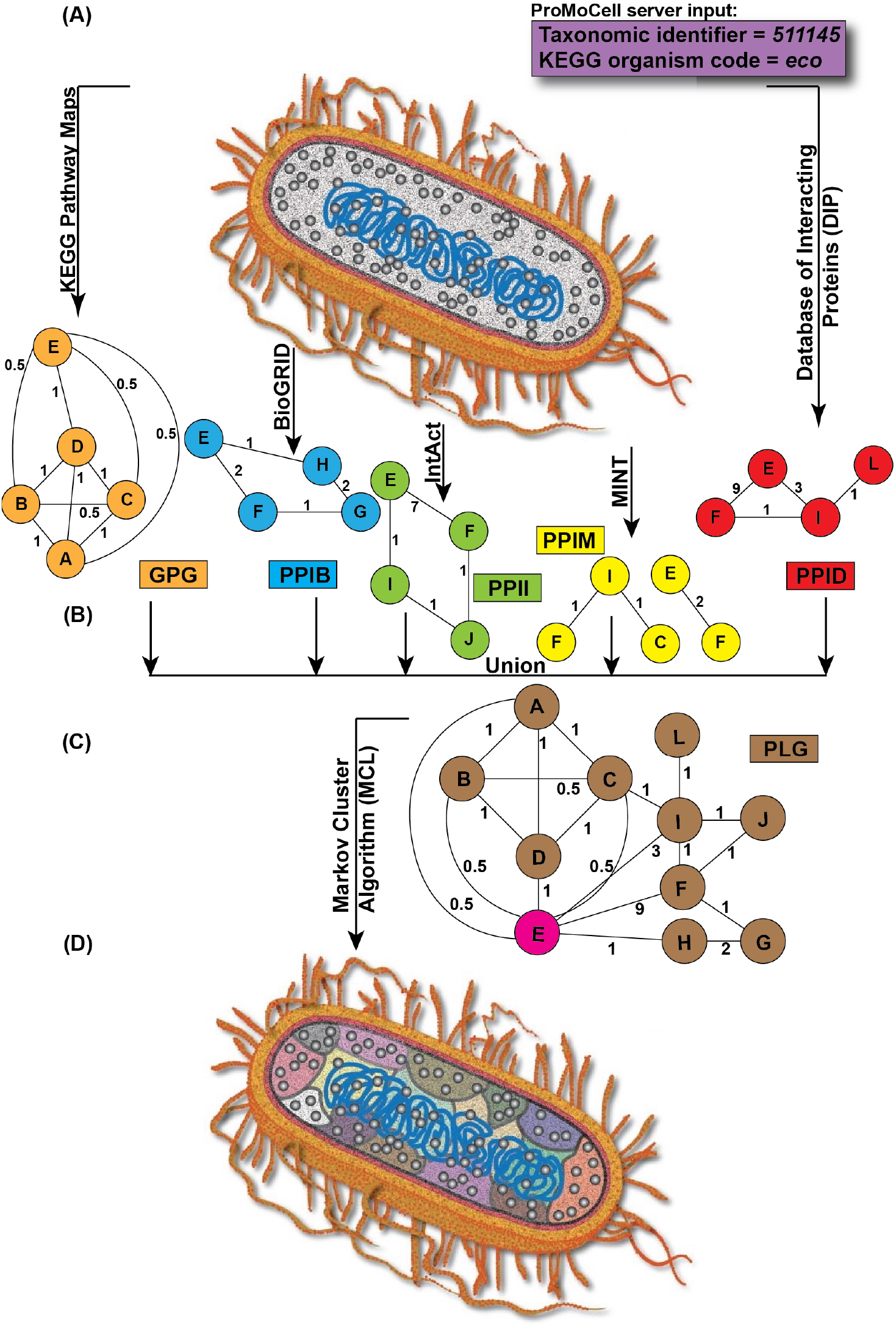
ProFuMCell methodology demonstration. (A) A schematic diagram of *Escherichia coli* (*E. coli*) cell whose cytoplasmic proteins are represented by grey spheres. (B) Construction of GPG, PPID, PPII, PPIM, and PPIB by utilizing existing knowledge from KEGG, DIP, IntAct, MINT, and BioGRID, respectively. (C) Assembling to PLG. (D) Schematic representation of spatially localized functional modules (colored regions in the cytoplasm of *E. coli*) as a result of clustering PLG using ProFuMCell.

### The database

One of the lacunas of ProFuMCell is its computation time which based on the input organism may take few hours. Therefore, we precompute the organism and store that information in a database for single-click download without any waiting time. Again, it shall reduce server repetitive computation on the same data at the server end. Neverthelss, ProFuMCell will be useful for unavailable organism or if user wish to recompute it afresh.

ProModb is a repository that documents computationally generated whole-cell proteinprotein interaction network-based functional modules of an organism using ProFuMCell. This database provides the users with a comprehensive and an integrated tool for an efficient browsing and extraction of the protein-protein interaction networks and the functional modules constructed by gathering experimental data from several interaction databases and pathway maps. Beyond indexing the functional modules and the protein-protein interaction networks, the ProModb is useful for understanding protein functions, studying the properties and topology of the protein interaction networks, examining the evolution of the protein-protein interactions and the individual proteins. ProModb is also useful for predicting the functions for the previously uncharacterized proteins, detecting novel pathways, elucidating the molecular basis of diseases thereby inspiring us in developing methods for the prevention, diagnosis, and treatment of the diseases, and identifying novel disease genes. A detailed schematic representation of the database structure is shown in Figure 2.

**Figure 2:**
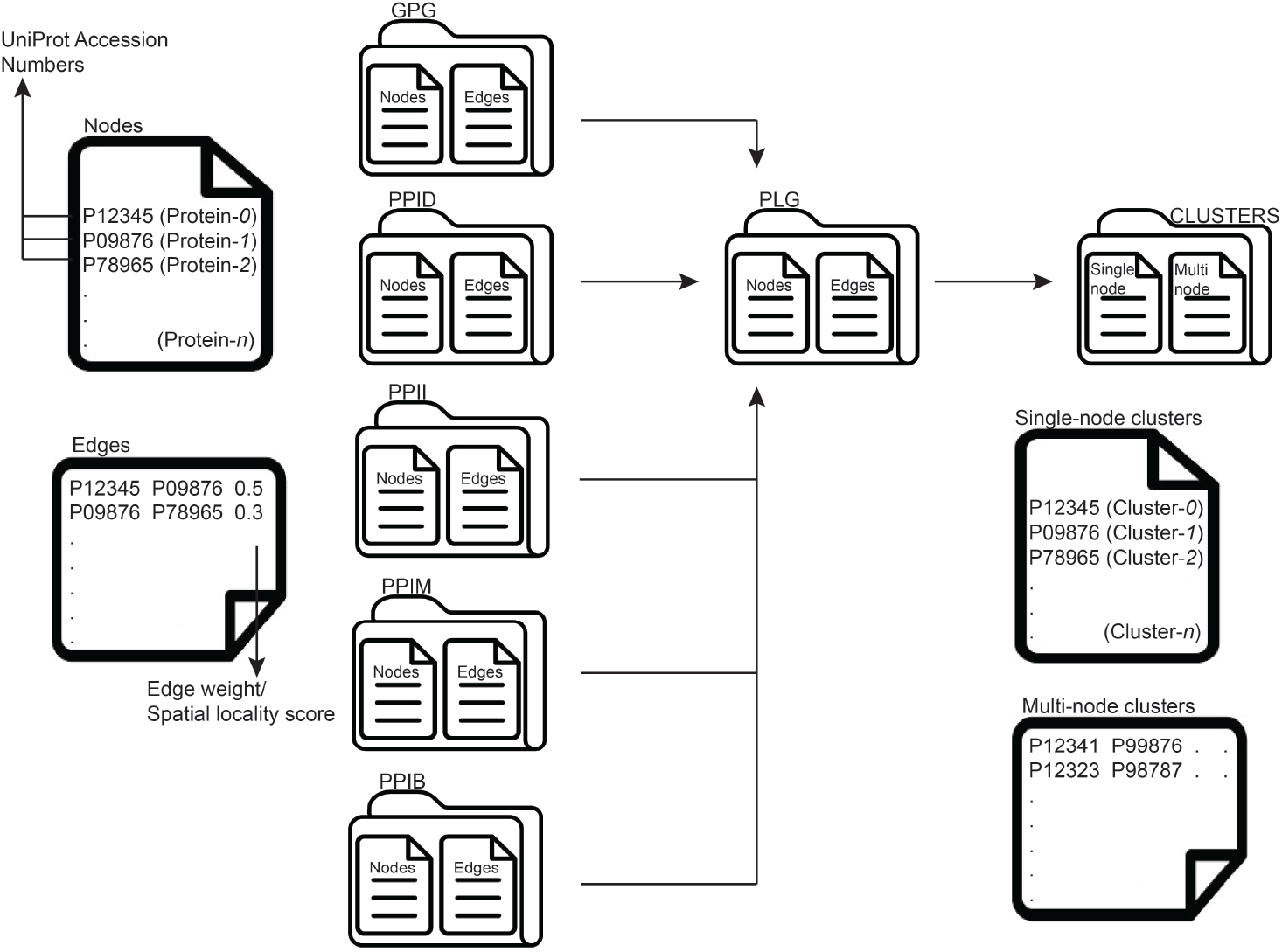
The structure of ProModb. The graphs including GPG, PPID, PPIM, PPII, PPIB, and PLG are present in their respective folders. Cluster/functional module information is present in a separate folder entitled as clusters. All are TSV files which can be easily downloaded, loaded into, and visualized using Cytoscape.

ProModb contemplates the experimentally verified actual physical interactions listed under the DIP, Molecular Interaction database, IntAct, and BioGRID. The database also considers the pathway information available in the KEGG pathway database. Such pathway maps do not represent actual physical interaction among the proteins, but are beneficial towards representing the potential protein-protein interactions. Information on the PPID, PPIM, PPII, PPIB, PLG, and functional modules, generated by the network-based zoning approach, ^8^ are integrated together and provided to the user in a simple user-friendly environment by ProModb. Despite the existence of several organism specific databases, the ProModb aims to include spatial locality-based protein interaction networks and functional modules for many organisms. Currently, ProModb supports 607 organisms belonging to the KEGG genome or the collection of KEGG organisms, including 483 eukaryotes and 124 bacteria.

For each organism, data in the ProModb is organized in the form of seven folders named as GPG, PPID, PPII, PPIM, PPIB, PLG, and clusters. The Protein Interaction Networks (node and edge information) corresponding to GPG, PPII, PPIM, PPID, PPIB, and PLG are present in their corresponding folders in the form of text files. The nodes of the PINs are the proteins which are represented by their corresponding Universal Protein Resource (UniProt)^16^ accession numbers. The edge weights of the interactions in the networks denote the respective spatial locality scores. If no experimental biological knowledge is available for a particular organism, then blank text files will be embedded into the database for the corresponding organism. All the text files used for the data storage in the ProModb follow TSV (Tab Separated Values) format, and hence can be easily downloaded and visualized using Cytoscape.^17^

Currently, ProModb contains organisms data belonging to the KEGG genome. Functional modules and the protein-protein interaction networks are thoroughly tested for their correctness. ProModb can be searched in a variety of ways by using one or a combination of the fields like organism name, taxonomic identifier, and the KEGG organism code.

### Server interface

#### ProModb

ProModb provides a user-friendly and straightforward interface for searching the database for a particular organism and viewing its corresponding PLG statistics or downloading the respective data from the server. ProModb interface, shown in Figure S3(A), lists the organisms along with their corresponding taxonomic identifiers, KEGG organism codes, categories of organisms, and the dates on which ProModb data are generated and uploaded to the database server. By default, the server displays 10 entries per page in lexicographical sorting of all the ProModb fields - organism, tax id, org code, category, data, and date, in both ascending and descending order. A separate search button allows the user to query for an organism from ProModb. For a particular organism, its corresponding date attribute represents the date on which its PLG-based protein modules are generated. The output and data attributes of ProModb provide data visualizations and a link to download all the generated data of an organism for further analysis. The downloaded folder enlists the functional modules along with their constituent proteins which are represented by their corresponding UniProt^16^ accession numbers. Additionally, PLG and all the constructed PPI networks are also available. All of these networks and the functional modules are Cytoscape compatible and hence, can be downloaded for visualization using Cytoscape.^17^

#### ProFuMCell

##### **I**nput

ProFuMCell provides a user-friendly and straightforward interface (Figure S1, S2) for job submission and result interpretation. The user needs to provide only the KEGG organism code (three- or four-letter code) and the taxonomic identifier (one to seven digits) that are the unique identification of an organism assigned by the KEGG genome^10^ and the National Center for Biotechnology Information (NCBI) respectively. ProFuMCell is exceptionally robust as it is capable of extracting interaction and pathway information of an organism from a diverse set of online resources by utilizing only the above-mentioned two simple input parameters that accurately characterize the organism. ProFuMCell constructs the interaction network for the whole-cell of an input organism by exploiting the experimentally verified data from KEGG pathway maps^10^ and protein-protein interaction databases like DIP,^11^ IntAct,^12^ MINT,^13^ and BioGRID^14^ as shown in Figure 1(A,B) and completes rest of the task in the ProFuMCell server itself. The server has got real-time web scraping characteristics and hence, the user will always get the latest information as per the pathway and interaction databases. The entire process is computation-intensive and may require few hours to complete based on the input organism and our server’s workload. We encourage the users to furnish his/her e-mail address (optional) for job notifications (submission and completion) else save the result page link. In case of invalid input, ProFuMCell will notify the mistake, and will discard the job.

##### **O**utput

ProFuMCell result page enlists the functional modules along with their constituent proteins as shown in Figures S3(B), S7, and S8. Additionally, PLG and all the constructed PPI networks are also available on the result page as depicted in Figures S4, S5, and S6. Each line of the tab delimited files storing the PPI networks (GPG, PPID, PPII, PPIM, PPIB) and the PLG represents each edge of the corresponding network. Thus, each line contains information on the nodes/proteins connected by an edge which are represented by their corresponding UniProt accession numbers. The edge weight denotes the spatial locality score which is represented by real numbers. Higher the weight of the edge connecting two proteins, lesser is the distance between them. ProFuMCell also provides the users with information on the single-node clusters or the self-interacting proteins whose two or more copies can interact with each other. The functional modules or the multi-node clusters generated by the web server consists of the spatially and functionally localized protein modules. Each line of the multi-node clusters file denotes a single cluster by listing its constituent proteins in the form of UniProt accession numbers. All of these networks and the functional modules can be downloaded for further analysis and easy visualization using Cytoscape software. ^17^

## Results and Discussion

ProFuMCell is wholly synchronized with the KEGG database^10^ and allows the users to generate functional modules for any organism belonging to the KEGG genome using its real-time web scraping characteristics. Hence, the server will host as many organisms as is maintained by KEGG database. Currently, ProFuMCell supports 7179 organisms, including 541 eukaryotes, 5955 bacteria, 334 archaea, and 349 viruses (Table 1). This support will go on increasing as and when new organism will be added into the KEGG genome and the corresponding protein interaction data will be incorporated into the PPI databases.

**Table 1:**
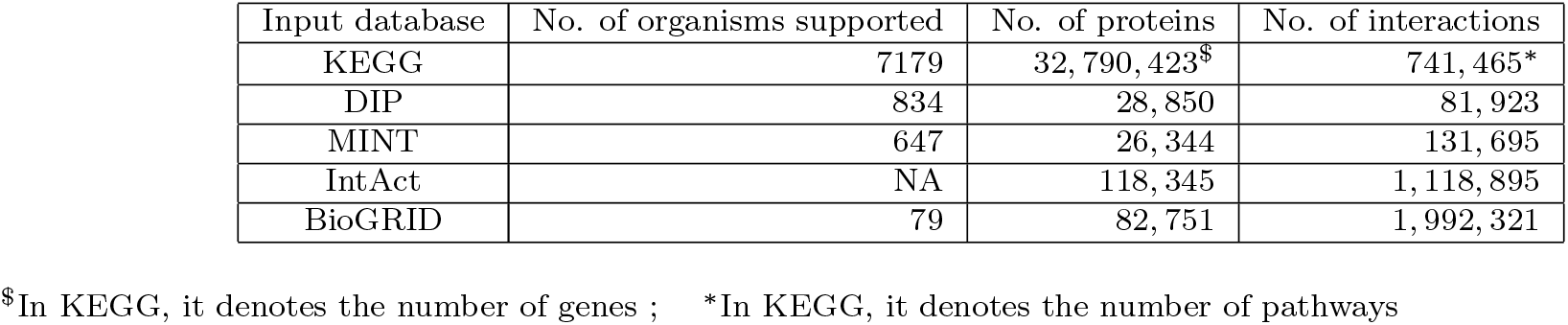
ProFuMCell statistics as of November 2, 2020. The table shows total number of organisms, proteins, and PPIs supported by ProFuMCell. This support will go on increasing along with the inclusion of new data into KEGG database, DIP, MINT, IntAct, and BioGRID.

### **C**urrent state of ProModb

ProModb contains data for 55 mammals, 20 birds, 10 reptiles, 3 amphibians, 32 fishes, 1 lancelet, 1 ascidian, 2 echinoderms, 1 hemichordate, 56 insects, 1 crustacean, 6 nematodes, 1 annelid, 5 mollusks, 1 brachiopoda, 4 flatworms, 6 cnidarians, 1 placozoan, 1 poriferan, 102 plants, 124 fungi, 50 protists, and 124 bacteria. Table 2 presents a brief analysis of the data stored in the server for different organisms. Data generation for the archaea and viruses are underway and accordingly the database is updated regularly. Thus, the support provided by the ProModb will go on increasing along with the introduction of newly discovered organisms to the KEGG genome. The largest Protein Interaction Network in ProModb is that of human, which contains a total number of 41550 nodes and 8943744 edges. The implementation details of ProFuMCell and ProModb are provided in the Supplementary Section S6.

**Table 2:** A brief analysis of the data stored in the ProModb server for organisms belonging to different categories.

| Organism | Category | #PLG Nodes | #PLG Edges | #Single-node clusters | #Multi-node clusters |
| --- | --- | --- | --- | --- | --- |
| *Xenopus laevis* | Amphibian | 6878 | 1434542 | 7 | 283 |
| *Helobdella robusta* | Annelid | 1729 | 93711 | 2 | 101 |
| *Ciona intestinalis* | Ascidian | 1462 | 62909 | 6 | 112 |
| *Escherichia coli ATCC 8739* | Bacteria | 289 | 1840 | 1 | 55 |
| *Anas platyrhynchos* | Bird | 250 | 1058 | 4 | 57 |
| *Hydra vulgaris* | Cnidarian | 384 | 2240 | 4 | 80 |
| *Danio rerio* | Fish | 4194 | 686180 | 28 | 212 |
| *Anopheles gambiae* | Insect | 1496 | 60415 | 1 | 99 |
| *Bos taurus* | Mammal | 5181 | 1623913 | 22 | 269 |
| *Arabidopsis lyrata* | Plant | 3473 | 293976 | 1 | 116 |
| *Alligator mississippiensis* | Reptile | 1303 | 46391 | 6 | 128 |

#### Mapping pathways to protein-protein interactions

KEGG^10^ is one of the most widely used online resources for studying the molecular interactions and the biochemical reactions happening inside the cell of an organism. KEGG encodes the high-level functions and the biological utility information in the form of pathway maps which are molecular interaction/reaction network diagrams. Every map is manually drawn by an in-house software called KegSketch. The nodes or entities participating in the pathway maps are either genes or compounds. As a result, KEGG reaction network diagrams highlight three types of possible interactions including gene-gene, compound-compound, and gene-compound. Parsing these KEGG provided molecular reaction network diagrams and mapping the pathways to PPIs is a difficult task. For performing these, we design a parser and a mapper which are already parts of the ProFuMCell package and is freely available for the research community. After parsing the pathway data, ProFuMCell contemplates only the gene-gene interactions of KEGG data and use the UniProt database^16^ to map the gene identifiers to their corresponding protein ids. Thus, the gene-gene interactions are converted to their corresponding protein-protein interactions by the server. ProModb stores this pathway mapped PPI data in the form of Gene Pathway Graph (GPG) for 607 organisms. ProFuMCell can perform real-time mapping of pathway data to their corresponding PPI data for any organism belonging to the KEGG genome. These pathway information mapped to the PPI data provide us with a set of potential interactions to be experimentally validated and will be useful for future analysis and research.

#### Protein-protein interaction prediction

There has always been a significant interest among the Systems Biology community for the computationally predicted protein-protein interactions. This is mainly because the experimental PPI data coverage is still very limited and relatively noisy.^18^ Along with incorporating the available experimentally verified PPI data, ProFuMCell and ProModb compute spatial locality-based PPI predictions which are currently unavailable. The predictions are computed on a graph utilizing the existing experimental data. ProFuMCell constructs the graphs by combining PPID, PPII, PPIM, PPIB, GPG, and PLG information, where protein molecules are acted as nodes/vertices. The experimentally verified interactions are represented as direct edges in these graphs, whereas the indirect edges are constructed using the transitivity property of the direct edges. The edge weights denote the spatial locality scores i.e., higher the edge weight, lesser is the distance between them, and higher is the reliability of the corresponding interaction.

For validation, we identify the spatially localized densely connected sub-networks from the PLG, which we find to be functionally close too through a detailed Gene Ontology-based analysis as performed in. ^8^ These locality-based PPI predictions are independent of other PPI predicting computational approaches. Such predictions could be utilized for those proteins which are not well-covered by the experimental PPI databases or to shortlist a set of potential interactions to be experimentally validated. Recently, machine-learning and deep-learning-based approaches are widely utilized for various research domains in Bioinformatics. Such approaches are proven to function well with the availability of more and more data. Hence, in case of limited experimental PPI datasets, our server predicted PPIs can be utilized for providing volume to the positive or negative training dataset required for machine and deep learning-based PPI prediction methods.

#### Deciphering biological evolution

Research conducted in^19,20^ provides new evidence that the topology of the Protein Interaction Networks contains information about the evolutionary processes. Again, detailed analyses on the functional modules of budding yeast and fission yeast reveal conservation of the intramodule connections, whereas, substantially differing cross-talk between the modules results in biological evolution.^21^ Since 1999, it has been noted that the intra-module connections stay conserved whereas the inter-cluster connections change during evolution. ^22^ Based on these observations, we understand that traces of evolution are not only present at the level of the Protein-Protein Interactions, but are also very much present at the level of the inter-module interactions. So, there exists a fair chance of phylogeny reconstruction by analyzing the topology of only the cross-talk between the modules rather than focusing on the conserved intra-module associations. Again, module-based analysis will be less time consuming since it mainly relies on the analysis by considering the comprehensive network properties like the network proximity and modularity rather than contemplating the individual interactions of the PPI network. ^23^ Therefore, detailed topological analyses of the Protein Locality Graphs of different organisms and their corresponding functionally localized modules stored in the ProModb server may be useful for deciphering biological evolution among the species.

#### Parallel whole-cell simulation

Modular cell biology states that different cellular functions are accomplished by different modules consisting of multiple interacting molecules. ^22^ These modules function within a particular compartment of the cell and their component molecules rarely traverse from its assigned region to other cellular compartments. ^1^ Accordingly, the spatially localized protein modules generated by ProFuMCell and stored by ProModb will be allotted certain subcellular compartments within which they will perform the cellular functions. Thus, the whole-cell simulation problem can be broken down into sub-cell simulation where each sub-cell along with its assigned module may be simulated on a separate computing unit of the High Performance Computing (HPC) systems as is demonstrated by Das et al.^8,9^ Granularity analysis of the PLG clusters of an organism generated by ProFuMCell will guide us in selecting a suitable HPC architecture for performing its whole-cell simulation. For example, there are 122 PLG multi-node clusters of an *Escherichia coli* cell, and four out of these 122 clusters consist of 1693, 771, 585, and 125 proteins. Rest of the clusters’ granularity vary between 2 and 23. Accordingly, the four enormous clusters are valid for a GPU simulation, whereas, the remaining clusters should be simulated in CPU since allocating small clusters to GPU will degrade the overall performance. Therefore, CPU-GPU systems or hybrid HPC architecture is most suitable for achieving the highest performance for the parallel whole-cell simulation of *E. coli*.^8,9^

## Case study

We consider *E. coli K12* strain for demonstrating the case study in this paper. *E. coli K12* is represented using the taxonomic identifier, ‘511145’ and the KEGG organism code, ‘eco’. Based on these two sole input parameters, ProFuMCell firstly extracts real-time pathway and protein interaction information from KEGG, DIP, MINT, IntAct, BioGRID and constructs GPG, PPID, PPIM, PPII, PPIB, respectively. From the extracted experimental knowledge, ProFuMCell computes spatial localities of the interacting proteins which are allocated as the edge weights in the constructed graphs. Finally, the web server merges all of these constituent networks into a single Protein Locality Graph. Each of GPG, PPID, PPIM, PPII, PPIB, and PLG consist of 1107 (43792), 2924 (12246), 258 (488), 3322 (20483), 139 (129), 3748 (64146) proteins (edges representing protein-protein interactions), respectively. ProFuMCell then applies Markov Cluster Algorithm to identify the densely connected protein clusters from the PLG. For *E. coli*, ProFuMCell identifies 188 clusters or functional modules which have been validated by a detailed Gene Ontology-based association performed in an earlier study. ^8^ ProFuMCell computed all these *E. coli* data has already been uploaded to ProModb which can be easily downloaded by searching the database service using organism name or taxonomic identifier or organism code. However, ProFuMCell should be executed whenever the user prefers to access up-to-date real-time computed data.

## Data and Software Availability

Publicly available ProFuMCell (https://cosmos.iitkgp.ac.in/ProFuMCell) and ProModb (https://cosmos.iitkgp.ac.in/ProModb) are free for all the users, and there is no login requirement. The services are effective for analyzing and simulating whole-cell.

## Conclusion

Two services, ProFuMCell and ProModb, are described for generating the Protein Locality Graph and the spatially localized functional modules of the cells of the organisms. ProFuMCell provides users with the real-time web scraping facilities and thereby, allowing them to generate the most recent and up to date PLG of the organisms. ProModb, on the other hand, stores the pre-generated data for a few selected organisms. We update ProModb on a regular basis to increase the support of organisms. Data provided by both the web services are beneficial for mapping pathways to PPIs, spatial locality-based prediction of PPIs, determining evolution, and for parallel whole-cell simulation. In the future, we hope to collect information on organism specific functional modules from databases (if available) and literature, link them with ProModb, and perform a comparative study between them and the ProModb provided modules.

## Declarations

### Availability of data and material

Publicly available ProFuMCell (https://cosmos.iitkgp.ac.in/ProFuMCell) and ProModb (https://cosmos.iitkgp.ac.in/ProModb) are free for all the users, and there is no login requirement.

### Code availability

N/A

### Authors’ contributions

B.D. and P.M. conceived the problem; B.D. designed and implemented the servers; B.D. analyzed the data; B.D. and P.M. wrote the paper.

