## Supplementary file for "ProFuMCell and ProModb: web services for analyzing interaction-based functionally localized protein modules in a cell"

#### Contents

**S1 ProMoCell** **2**

**S2 Initialization** **2**

**S3 Processing** **2**

**S4 Production** **3**

**S5 Interface** **4**

**S6 Implementation** **4**

#### S1 ProMoCell

ProMoCell (*Pro*tein interaction based functional *Mo*dules of the *Cell* of an organism) is a web server to determine the functional modules of the cell of an organism. ProMoCell is designed following our recent work [1] on the network-based zoning approach for whole-cell simulation. ProMoCell can identify clusters of spatially localized proteins from the Protein-Protein Interaction (PPI) network of the whole-cell of an organism. A thorough Gene-Ontology [2] based analysis of the clusters revealed that the spatially localized clusters are also in functional proximity and hence can be termed as the functional modules of the cell of the organism [1]. ProMoCell is a single-click web service, and it is straightforward, user-friendly and easy to use. Presumably, no other web service like ProMoCell exists till date.

#### S2 Initialization

The only user input required by ProMoCell is the KEGG organism code (three- or four-letter code) and the organism taxonomic identifier (one to seven digits) that is the unique identification of an organism assigned by the KEGG genome and the NCBI respectively. The server has got real-time web scraping characteristics and based on the input parameters, it constructs the interaction network for the whole-cell of an organism utilizing the experimentally verified data from various sources like KEGG pathway maps [3] and protein-protein interaction (PPI) databases like DIP [4], IntAct [5], MINT [6] and BioGRID [7].

#### S3 Processing

ProMoCell implements the network-based zoning approach published in [1]. For the given organism, ProMoCell starts by creating interaction networks from each of the above-mentioned databases. Gene Pathway Graph (GPG) is constructed by studying the gene-gene interactions of the KEGG pathway maps. Since genes undergo transcription and translation to synthesize proteins, so equivalently we focused on protein-protein interactions rather than gene-gene interactions. The set of proteins represents the vertices of GPG. Corresponding to each experimentally verified interaction as per KEGG database there is an edge between the corresponding proteins. These edges are unit weighted and are termed as DiEdge. Following the observation that two interacting proteins must be in proximity to each other, we added additional edges between those pairs of proteins whose interactions are not experimentally verified, and their weights are allocated as the inverse of the shortest DiEdge path weight computed using Dijkstra's algorithm [8]. Therefore, edge weight denotes the spatial locality score (higher the edge weight, lesser is the distance between them) between the corresponding proteins.

Next ProMoCell constructs PPI graphs namely PPID, PPIM, PPII, and PPIB by collecting the PPI data corresponding to the specific organism from four different interaction databases viz, Database of Interacting Proteins (DIP), Molecular Interaction Database (MINT), IntAct, and BioGRID respectively. Similar to GPG, unit weighted DiEdges are constructed between the pairs of proteins whose interactions are experimentally verified as per DIP (PPID graph), MINT (PPIM graph), IntAct (PPII graph), and BioGRID (PPIB graph). Then the edge weights are updated using the density of interactions to identify the weighted dense subgraphs of the PPI. The modified edge weight was directly proportional to the edge cut of the interaction pair which was computed by the max-flow-min-cut theorem [9]. Then ProMoCell assembles the Protein Locality Graph (PLG) by merging the GPG, PPID, PPII, PPIM, and PPIB. The vertices of PLG

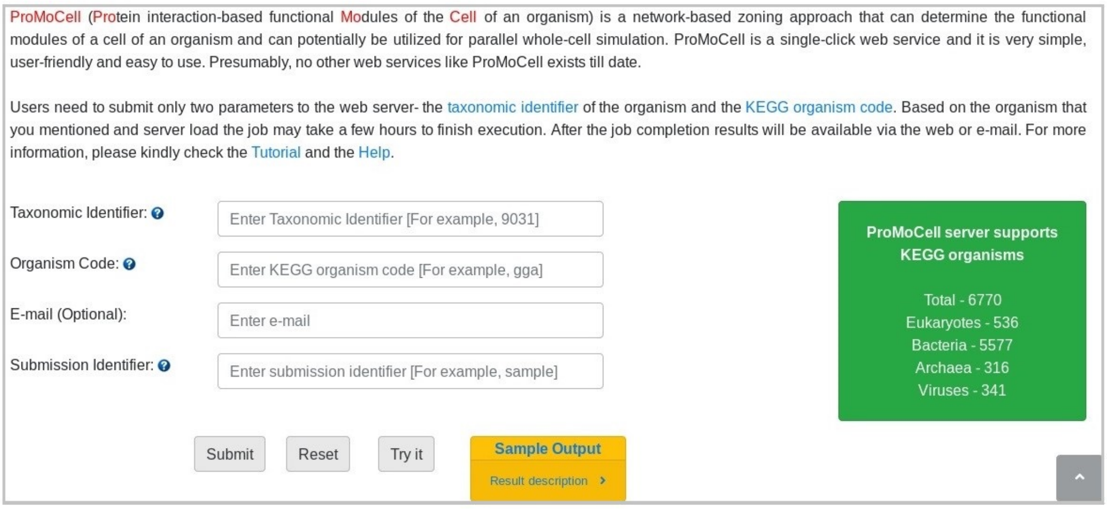


**Figure S1:** ProMoCell server interface.

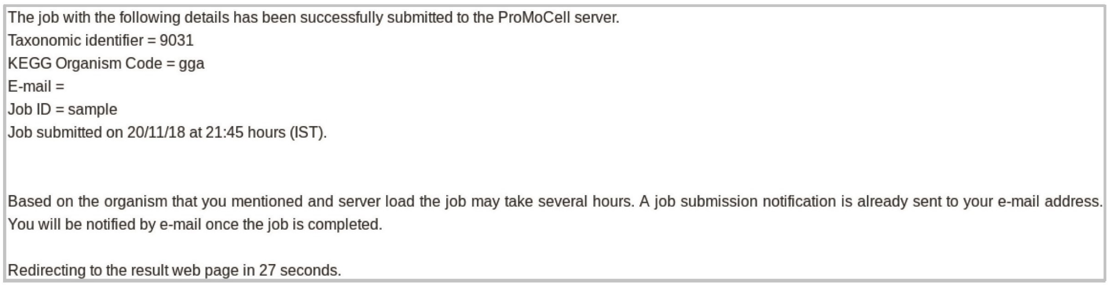


**Figure S2:** ProMoCell job submission webpage.

represent the union of the vertices of the GPG and all the PPI graphs. The edge weights of PLG is decided as the maximum of the edge weights of the component graphs. Finally, the Markov Cluster Algorithm [10] is applied to the PLG to generate the spatially localized clusters from the densely connected sub-graphs of the PLG [1]. The sole parameter guiding MCL is the inflation parameter which regulates the cluster granularity, and there is no fixed rule to determine the correct value of the parameter which mainly depends on the nature of the input graph. Mostly PLG is a disconnected graph consisting of one or multiple single-node components which contradict our objective of uniform cellular division. So, we experiment with different values of inflation parameters and select the highest possible one which does not generate any new single-node cluster.

### S4 Production

ProMoCell result page enlists the functional modules along with their constituent proteins which are represented by their corresponding UniProt accession numbers. Additionally, PLG and all PPI networks are also available on the result page (Supplementary Figure S1-4). The functional modules generated by the web server can be utilized for determining the evolutionary orthology signal [11]. Furthermore, a detailed comparative analysis can be performed on the functional modules of the healthy and diseased cells which can eventually aid us in identifying the source or cause of diseases in the cell. In other words, proper knowledge of the functional modules can assist in the diagnosis, prevention, and treatment of deadly diseases like cancer [12]. The result page also provides a link to download all the generated data for further analysis.

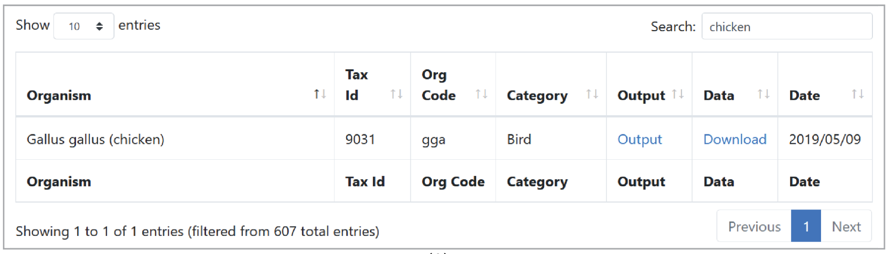


(A)

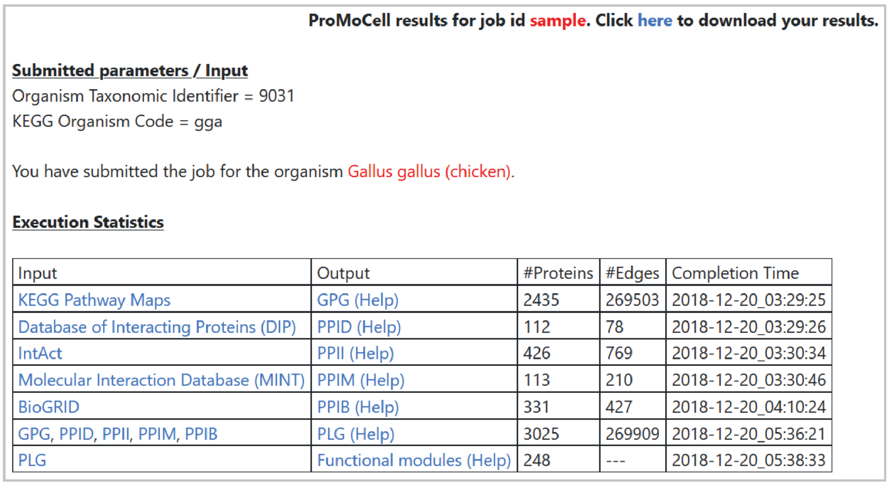


(B)

**Figure S3:** (A) ProModb server interface displaying the search result for the query organism, chicken along with its corresponding taxonomic identifier, KEGG organism code, category, and the date on which ProModb data are generated and uploaded to the database server. (B) The result page of ProMoCell for the input organism *Gallus gallus* (chicken) along with the Table of the execution statistics.

#### S5 Interface

The only user input required by ProMoCell is the KEGG organism code (three- or four-letter code) and the organism taxonomic identifier (one to seven digits) that is the unique identification of an organism assigned by the KEGG genome and the NCBI respectively. The ProMoCell server interface and the job submission webpage are shown in Supplementary Figures S1 and S2, respectively. The entire process is computation intensive and may require few hours to complete based on the input and the server load. We encourage the users to furnish his/her email address for automatic email notifications regarding job status (submission and completion). Nonetheless, contact information is optional, and a user may wish to save or bookmark the link of the result page without providing an email address. The server, connected with a high-performance computing cluster, is capable of handling multiple tasks in parallel. All job identifiers are randomly generated, and hence each of them is entirely confidential in the server. ProMoCell is free and open to all users, and there is no login requirement. For more information on the server methodology and its usage, kindly refer to `https://cosmos.iitkgp.ac.in/ProMoCell/tutorial.php` and `https://cosmos.iitkgp.ac.in/ProMoCell/help.php`.

#### S6 Implementation

ProMoCell and ProModb, both are designed using PHP v7.0.32 (front end) on a Linux platform, along with Java v9 and Python v3.9.0 (backend). The server, connected with a high-performance computing cluster, is capable of handling multiple tasks in parallel. All job identifiers are randomly generated, and hence each of them is entirely confidential

**Edges of GPG**

| | $Node_1$ | $Node_2$ | Weight |
|---|---|---|---|
| $Edge_1$ | F1ND09 | Q90978 | 0.33 |
| $Edge_2$ | Q27IP4 | Q9PU44 | 0.33 |
| $Edge_3$ | E1C3V3 | E1C795 | 0.33 |
| . | . | . | . |
| . | . | . | . |
| $Edge_{269503}$ | . | . | . |

**Nodes of GPG**

| $Node_1$ | $Node_2$ | $Node_3$ | . | . | $Node_{2435}$ |
|---|---|---|---|---|---|
| R4GKI4 | Q5ZLA5 | Q5ZLA7 | . | . | . |

**Edges of PPID**

| | $Node_1$ | $Node_2$ | Weight |
|---|---|---|---|
| $Edge_1$ | P09572 | P08251 | 1.0 |
| $Edge_2$ | P00523 | Q05860 | 2.0 |
| $Edge_3$ | Q05860 | Q05876 | 2.0 |
| . | . | . | . |
| . | . | . | . |
| $Edge_{78}$ | . | . | . |

**Nodes of PPID**

| $Node_1$ | $Node_2$ | $Node_3$ | . | . | $Node_{112}$ |
|---|---|---|---|---|---|
| B0I564 | D2YW45 | D9N195 | . | . | . |

**Figure S4:** Edges and nodes of GPG and PPID as displayed in the Result page of the ProMoCell for the input organism Gallus gallus (chicken).

**Edges of PPII**

| | $Node_1$ | $Node_2$ | Weight |
|---|---|---|---|
| $Edge_1$ | P31946 | P07228 | 3.0 |
| $Edge_2$ | P07228 | P31946 | 3.0 |
| $Edge_3$ | P31946 | P07228 | 3.0 |
| . | . | . | . |
| . | . | . | . |
| $Edge_{769}$ | . | . | . |

**Nodes of PPII**

| $Node_1$ | $Node_2$ | $Node_3$ | . | . | $Node_{426}$ |
|---|---|---|---|---|---|
| A0A1D5P7R1 | A4VAR9 | A7MB62 | . | . | . |

**Edges of PPIM**

| | $Node_1$ | $Node_2$ | Weight |
|---|---|---|---|
| $Edge_1$ | P01110 | P52162 | 4.0 |
| $Edge_2$ | P01110 | P52162-2 | 4.0 |
| $Edge_3$ | P01109 | P52162-2 | 4.0 |
| . | . | . | . |
| . | . | . | . |
| $Edge_{210}$ | . | . | . |

**Nodes of PPIM**

| $Node_1$ | $Node_2$ | $Node_3$ | . | . | $Node_{113}$ |
|---|---|---|---|---|---|
| P18433 | Q90574 | Q99LM3 | . | . | . |

**Figure S5:** Edges and nodes of PPII and PPIM as displayed in the Result page of the ProMoCell for the input organism Gallus gallus (chicken).

**Edges of PPIB**

| | Node$_1$ | Node$_2$ | Weight |
|---|---|---|---|
| Edge$_1$ | P62760 | P60706 | 1.0 |
| Edge$_2$ | P62764 | P60706 | 1.0 |
| Edge$_3$ | A0A0B4J1S3 | F1NBT9 | 1.0 |
| . | . | . | . |
| . | . | . | . |
| Edge$_{427}$ | . | . | . |

**Nodes of PPIB**

| Node$_1$ | Node$_2$ | Node$_3$ | . | . | Node$_{331}$ |
|---|---|---|---|---|---|
| A0A024QZJ8 | A0A024R597 | A0A024R9A9 | . | . | . |

**Edges of PLG**

| | Node$_1$ | Node$_2$ | Weight |
|---|---|---|---|
| Edge$_1$ | A0A024QZJ8 | F1NAP2 | 1.0 |
| Edge$_2$ | A0A024R597 | A0A1D5PC18 | 3.0 |
| Edge$_3$ | A0A024R597 | P0C1H3 | 2.0 |
| . | . | . | . |
| . | . | . | . |
| Edge$_{269909}$ | . | . | . |

**Nodes of PLG**

| Node$_1$ | Node$_2$ | Node$_3$ | . | . | Node$_{3025}$ |
|---|---|---|---|---|---|
| P22770 | A0A1D5PYQ3 | F1NYJ1 | . | . | . |

**Figure S6:** Edges and nodes of PPIB and PLG as displayed in the Result page of the ProMoCell for the input organism Gallus gallus (chicken).

**Single-Node Clusters of PLG**

| Node$_1$ | Node$_2$ | Node$_3$ | . | . | Node$_{15}$ |
|---|---|---|---|---|---|
| A0A1D5P5J3 | A0A1D5PQ40 | D2YW45 | . | . | . |

**Figure S7:** ProMoCell result of the single-node clusters for the cell of Gallus gallus (chicken) (the input organism).

**Multi-Node Clusters of PLG / Functional Modules of the cell of the specified organism**

| | Node$_1$ | Node$_2$ | Node$_3$ | . | . | Node$_{1054}$ |
|---|---|---|---|---|---|---|
| Cluster$_1$ | A0A060PJH4 | A0A0K0PVD8 | . | . | . | . |

| | Node$_1$ | Node$_2$ | Node$_3$ | . | . | Node$_{225}$ |
|---|---|---|---|---|---|---|
| Cluster$_2$ | P28701 | F1NHH9 | . | . | . | . |

| | Node$_1$ | Node$_2$ | Node$_3$ | . | . | Node$_{198}$ |
|---|---|---|---|---|---|---|
| Cluster$_3$ | A0A1D5P0J0 | A0A1D5PGV0 | . | . | . | . |

.
.

| | Node$_1$ | Node$_2$ |
|---|---|---|
| Cluster$_{248}$ | . | . |

**Figure S8:** ProMoCell result of the functional modules (multi-node clusters) for the cell of Gallus gallus (chicken) (the input organism).

in the server. Both the web services are free and open to all users, and there is no login requirement. Further information on the server methodology and its usage is available at the tutorial page (`https://cosmos.iitkgp.ac.in/ProMoCell/tutorial.php`), and at the help pages (`https://cosmos.iitkgp.ac.in/ProMoCell/help.php`), (`https://cosmos.iitkgp.ac.in/ProModb/help.php`).
